# An MST-based assay reveals new binding preferences of IFIT1 for canonically and non-canonically capped RNAs

**DOI:** 10.1101/2024.05.10.593534

**Authors:** Tomasz Spiewla, Katarzyna Grab, Anais Depaix, Kamil Ziemkiewicz, Marcin Warminski, Jacek Jemielity, Joanna Kowalska

**Affiliations:** Division of Biophysics, Institute of Experimental Physics, Faculty of Physics, University of Warsaw, Pasteura 5, 02-093 Warsaw; Doctoral School of Exact and Natural Sciences, University of Warsaw, Zwirki i Wigury 93, 02-089 Warsaw; Centre of New Technologies, University of Warsaw, Banacha 2c, 02-097 Warsaw

## Abstract

IFIT proteins (interferon-induced proteins with tetratricopeptide repeats) are key components of the innate immune response that bind to viral and cellular RNA targets to inhibit viral translation and replication. The RNA target recognition is guided by molecular patterns, particularly at the RNA 5’ ends. IFIT1 preferably binds RNAs modified with the 7-methylguanosine (m^7^G) cap-0 structure, while RNAs with cap-1 structure are recognized with lower affinity. Less is known about the propensity of IFIT1 to recognize non-canonical RNA 5’ ends, including hypermethylated and non-canonical RNA caps. Deciphering the structure-function relationship for IFIT1-RNA interaction may improve understanding of cellular selection of IFIT targets and guide the design of exogenously delivered therapeutic RNAs, but requires high-throughput and robust analytical methods. Here, we report a biophysical assay for quick, direct, in-solution affinity assessment of differently capped RNAs with IFIT1. The procedure, which relies on measuring microscale thermophoresis (MST) of fluorescently labelled protein as a function of increasing ligand concentration, is applicable to various RNA lengths and sequences without the need for labelling or affinity tagging. Using the assay, we examined thirteen canonically and non-canonically 5’-capped RNAs, revealing new binding preferences of IFIT1. The 5’ terminal m^6^A mark in the m^7^G cap had a protective function against IFIT1, which was additive with the effect observed for the 2’-O position (m^6^A_m_ cap-1). In contrast, an increased affinity for IFIT1 was observed for several non-canonical caps, including trimethylguanosine (TMG), unmethylated (G), and flavin-adenine dinucleotide (FAD) caps. The results suggest new potential cellular targets of IFIT1 and may contribute to broadening the knowledge on the mechanisms of the innate immune response as well as the more effective design of chemically modified mRNAs.

## INTRODUCTION

The innate immune response against foreign nucleic acids constitutes the initial cellular defense mechanism against bacteria and viruses (Kawasaki and Kawai 2019, Ghoreshi et al. 2022). In order to respond to viral infections, host cells need to identify the threat and activate efficient antiviral defense. This defense mechanism is set in motion in higher eukaryotes upon recognizing foreign pathogen-associated molecular patterns (PAMPs), such as unique viral RNA fragments, uncapped RNAs, and double-stranded RNAs. These viral PAMPs are identified by specific cellular pattern-recognition receptors (PRRs) such as Toll-like receptors (TLR3, TLR7, and TLR8) and RIG-I-like receptors (RLR: RIG-I and MDA-5), which are present both in endosomes and the cytoplasm, specifically recognizing foreign nucleic acids. (Kumar et al. 2014, Imaizumi et al. 2021, Chen et al. 2021, Rehwinkel and Gack 2020, Dias, Sampaio and Rehwinkel 2019). The interaction between these receptors and PAMPs triggers a signaling cascade resulting in the expression of type I antiviral interferons (IFN-α and -β)(Franco, Chattopadhyay and Pan 2023). Secreted IFN-α and IFN-β then bind to type I IFN receptors (IFNARs), inducing the expression of numerous IFN-stimulated genes (ISGs) that encode proteins with regulatory and antiviral functions (Hyde et al. 2014, Pidugu et al. 2019a). Prominent members of this group are interferon-induced proteins with tetratricopeptide repeats (IFIT), which play pivotal roles in diverse biological mechanisms such as virus-induced translation inhibition, PAMP recognition, cell proliferation, replication, and signaling of double-stranded RNA (Fleith et al. 2018, Fitzgerald and Kagan 2020, Li and Wu 2021). IFIT proteins are recognized as immediate antiviral effectors during the innate immune response in humans and other mammals. While lacking enzymatic functions, they directly binding to non-self and cellular RNAs, and proteins to achieve their antiviral effects (Menachery et al. 2014, Hyde and Diamond 2015, Daugherty et al. 2016).

IFIT genes have been identified in numerous mammals, but the number and composition of the IFIT gene family can vary significantly from one species to another (Fensterl and Sen 2011). The human family consists of four canonical members namely IFIT1, IFIT2, IFIT3, and IFIT5 as well as uncharacterized IFIT1B and pseudogene IFIT1P1 (Pidugu et al. 2019a, Daugherty et al. 2016, Fensterl and Sen 2015). IFITs typically remain silent or are expressed at low levels under normal conditions. They are rapidly expressed upon viral infection, reaching concentrations 100 to 1000 times higher in the cytosol where they interact with foreign RNA (Fensterl and Sen 2015). IFITs consist of multiple repetitions of the tetratricopeptide repeat (TPR), which is a helix-turn-helix motif consisting of 34 amino acids. The TPR sequences are characterized by a degenerate pattern, with only 9 residues at specific positions exhibiting limited conservation. Consequently, IFIT proteins display significant structural variability and different specificities towards RNA targets that are mostly guided by the structure of RNA 5’ terminus (Szretter et al. 2012). Both IFIT1 and IFIT5 directly bind to RNA – IFIT5 primarily recognizing uncapped RNA (i.e. 5′-triphosphorylated), while IFIT1 binds to RNAs carrying a cap-0 structure at the 5’ end (i.e. lacking the 2’-O methylation at the first transcribed nucleotide). This prevents the hijacking of the host’s translational machinery by the virus and inhibits viral replication (John et al. 2017). The antiviral effect of IFIT1 is enhanced in the complex with IFIT3 (Abbas et al. 2013, Katibah et al. 2013, Johnson et al. 2018, Pidugu et al. 2019b). The RNA binding site of IFIT1 is composed of a highly positively charged pocket with a distinguished 5’-end binding site, enabling the binding of various 5’-capped RNA ligands (Yang et al. 2012), (Choi, Bowman and Jung 2018). While IFIT1 protein predominantly resides in the cytoplasm, research has demonstrated that approximately 10-15% of endogenous IFIT1 is present in the cell nucleus, with active shuttling of the protein occurring between the two compartments.

The IFIT1-capped RNA interaction has been already examined using various methods, such as primer extension inhibition (Fleith et al. 2018), electromobility shift assays (Habjan et al. 2013), surface plasmon resonance (Pichlmair et al. 2011), filter binding (Kumar et al. 2014), or bio-layer interferometry (Miedziak et al. 2020). In all the studies reported so far, IFIT1 displayed binding preference for cap-0 over cap-1 or uncapped RNAs (Abbas et al. 2017). These investigations consistently revealed a high affinity of IFIT1 for cap-0-RNA, with dissociation constant values (*K*_*D*_) ranging from 23 to 175 nM, irrespective of the RNA’s origin (viral or host) and sequence. Comparable studies have indicated that IFIT1 also binds to 5’-triphosphate and cap-1-RNAs, albeit with notably lower affinities (*K*_D_ ∼250 nM to >1 μM and K_D_ ∼450 to 710 nM, respectively) (Pichlmair et al. 2011, Katibah et al. 2014, Kimura et al. 2013, Miedziak et al. 2020). As concluded from primer extension assays, the critical elements for IFIT1 binding include the 5’ terminal mRNA cap residue and at least 4-5 nucleotides downstream of the cap structure (Chung et al. 1994, Kumar et al. 2014). These additional nucleotides likely enhance the stability of IFIT1-RNA complex, blocking the recruitment of additional ligands to the binding pocket or eliminating nonspecific interactions by inducing conformational changes in the protein (Zhang et al. 2014). Significantly less is known about the affinity of IFIT1 for other cap structures, such as the non-canonical caps found on some endogenous, viral, and bacterial RNAs or chemically modified caps that are being developed for modification of *in vitro* transcribed mRNAs for therapeutic purposes (Warminski et al. 2023). We envisaged that a simple, easily accessible, material- and time-saving method for quantitative assessment of cap-IFIT would greatly expedite studies in this direction.

As such, we report here a simple, rapid, and cost-effective assay that enables the determination of the binding affinity of variously capped RNAs for IFIT1 based on microscale thermophoresis (MST) phenomenon (Duhr and Braun 2006, Jerabek-Willemsen et al. 2014, Scheuermann et al. 2016, Tso et al. 2018). The assay enables precise measurement of *K*_D_ values between fluorescently labeled IFIT1 and RNAs of various lengths. Utilizing this assay, we evaluated a library of 13 RNAs containing different 5’-terminal structures (Figure 1). Our exploration included back-to-back comparison of all four m^7^G cap-0 variants incorporating adenine, guanine, cytosine, and uridine as the 5’-terminal nucleobase, and some methylated derivatives commonly found in human mRNA, such as 2’-*O*-methyl and N6-methyl on adenosine as the 5’-terminal nucleoside. We also investigated several non-canonical caps (NNCs), including nicotinamide-adenine dinucleotide (NAD) found on some RNA across all domains of life, an unmethylated “G” cap that is associated with aberrant mRNA capping, hypermethylated trimethylguanosine (TMG) cap present in snRNAs but also found on some Rev/RRE-dependent HIV-1 RNAs (Yedavalli and Jeang 2010, Chen et al. 2009, Julius and Yuzenkova 2017), UDP-Glucose cap, which is a relatively abundant non-canonical RNA cap, and flavin-adenine dinucleotide cap (FAD), which has been recently identified as the major molecule modifying the 5’ end of HCV RNA (Figure 1) (Sherwood et al. 2023, Wang et al. 2019). Moreover, we also investigated if the use of trinucleotide ligands can provide similar quantitative information regarding cap-IFIT interaction as the use of long-capped RNAs. The results revealed new 5’-terminal moieties that unexpectedly stabilize the interaction of capped RNA with human IFIT1, highlighting new potential cellular targets and underscoring the need for further investigation of the RNA cap-related aspects of the innate immune response.

**Figure 1.**
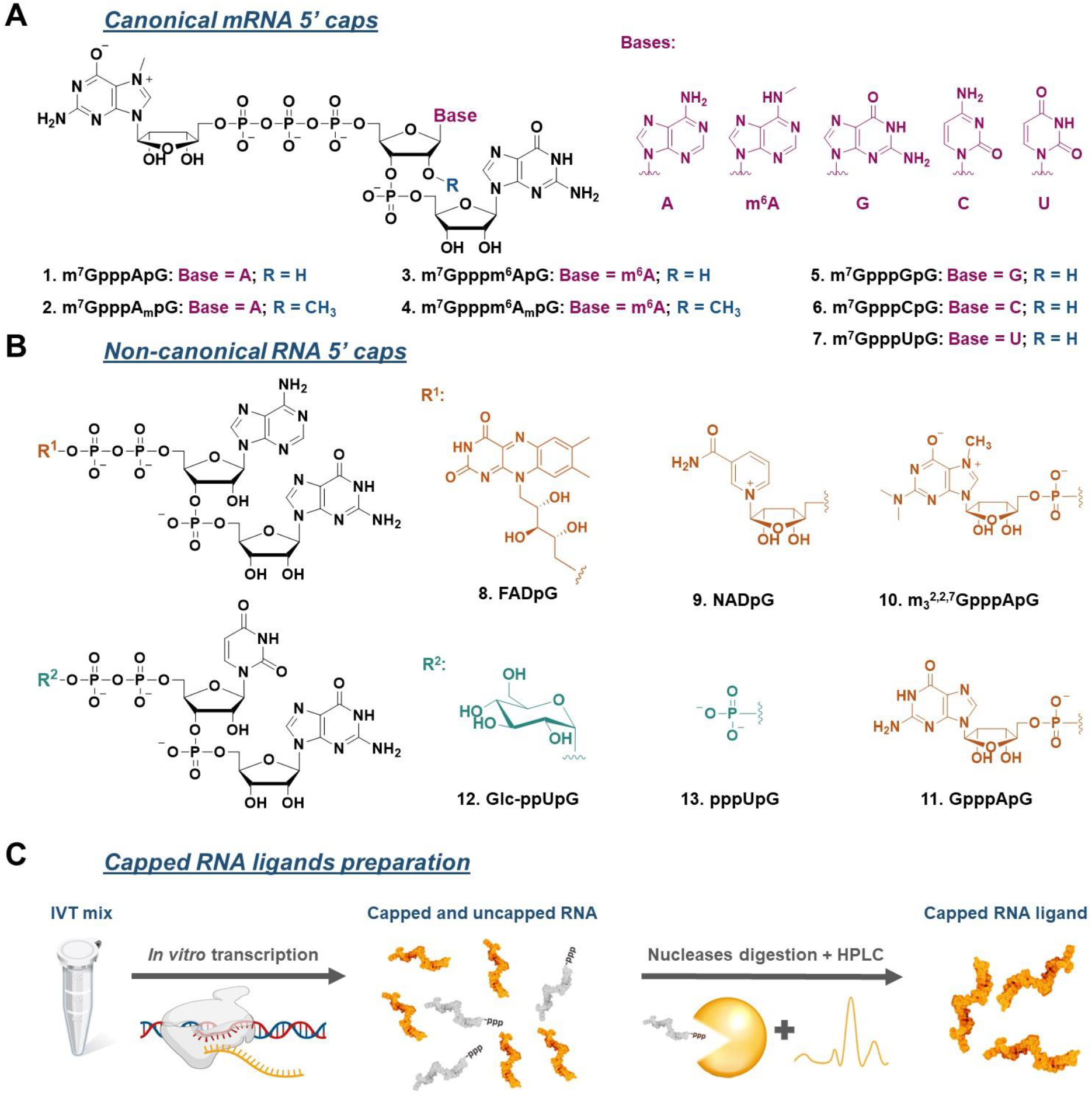
Structures of canonical and non-canonical RNA caps studied in this work. Structures of the A) canonical and B) non-canonical RNA 5’ cap analogs used for RNA preparation by *in vitro* transcription. C) The workflow for RNA preparation and purification leading to highly homogenous 5’ capped RNA.

## MATERIALS AND METHODS

### RNA Cap Analogs

Canonical 5’-cap analogs (m^7^GpppApG, m^7^GpppA_m_pG, m^7^GpppGpG, m^7^GpppCpG, m^7^GpppUpG, m^7^Gppp^m6^ApG, m^7^Gppp^m6^A_m_pG) were synthesized as previously described (Sikorski et al. 2020). Non-canonical 5’-cap analogs (NADpG, FADpG, Glucose-ppUpG) and uncapped pppUpG were synthesized as previously described (Depaix et al. 2022). 5’-cap analogs with modification in m^7^G structure (GpppApG and m_3_^2,2,7^GpppApG) were synthesized as previously described (Grab et al. in preparation).

### Synthesis of Capped RNA

The capped RNA ligands used in the study were synthesized by *in vitro* transcription (IVT) on the DNA template containing the T7 promoter ϕ6.5 (TAATACGACTCACTATAGGG). A typical transcription reaction (100 μL) was conducted for 1 hour at 37°C and contained the concentrations of nucleotides (NTPs, Thermo Fisher Scientific) and cap analogs as specified below. Transcription conditions for each class of cap analogs were optimized to ensure high incorporation of the analog at the RNA 5’ end (capping efficiency). For the cap analogs m^7^GpppApG, m^7^GpppA_m_pG, m7Gppp^m6^ApG and m7Gppp^m6^A_m_pG, the transcription mixture contained 5 mM concentrations of UTP/ATP/CTP, 4 mM GTP, and 10 mM trinucleotide cap analog. For the cap analogs m^7^GpppGpG, m^7^GpppCpG and m7GpppUpG, the transcription mixture contained 4 mM UTP/ATP/CTP, 2 mM GTP, and 8 mM trinucleotide cap analog. For all other analogs (NADpG, FADpG, Glc-ppUpG, m_3_^2,2,7^GpppApG, GpppApG and pppUpG) the transcription mixtures contained 4 mM UTP/ATP/CTP, 2 mM GTP and 12 mM trinucleotide/dinucleotide analog. The reaction mixtures were supplied with RNA polymerase buffer (Thermo Fisher Scientific), 20 mM MgCl_2_, 1U/μL RiboLock RNase Inhibitor (Thermo Fisher Scientific), 0.002 U/μL Inorganic Pyrophosphatase (Thermo Fisher Scientific), T7 RNA polymerase (20 U/μL Thermo Fisher Scientific) and 40 ng/μL annealed oligonucleotides (Genomed) as a DNA template. Three different sequences were used for RNA sequence optimization, presents as DNA oligonucleotides in supplementary, see Table S1. RNA I consisted of the following nucleotides: GGGACATTTGCTTCTGACACAACTGTGTTCACTAGCAA, RNA II – GGGAAAAAAAAAAAAAAAACATTTGCTTCTGACACA, and RNA III – GGGACATTTGCTTCTGACACAACTGTGTTCACTAGCAACCTCAAACAGACACCATGGTGCATCTGACTCCT. After 1 h incubation of *in vitro* transcription mix, 7 U of DNase I (Thermo Fisher Scientific) was added, followed by incubation for another 30 min at 37°C. To stop the reaction, 10 μl of 500 mM aqueous Na_2_EDTA solution (equimolar amount of EDTA respective to Mg^2+^) was added. The crude RNA was preliminarily purified using Monarch RNA Clean-up kit (New England Biolabs) according to the manufacturer’s protocol. Then a two-step enzymatic digestion using 5’ polyphosphatase (Lucigen) and Xrn-1 (New England Biolabs) enzymes was applied to all RNAs except pppUpG-RNA to remove uncapped RNA. The content (in μg) of uncapped RNA was estimated based on the amount of RNA material and the capping efficiency of the individual analogues. In the first step, the reaction with 5’ polyphosphatase (100 μL) was carried out for 1 h at 37°C and contained: digested RNA, 1 U/μL RiboLock RNase Inhibitor (Thermo Fisher Scientific), 5’ polyphosphatase buffer (Lucigen) and 5’ polyphosphatase (Lucigen), using 1 μL of enzyme per 5 μg of uncapped RNA. RNA samples were again purified with Monarch RNA Clean-up (New England Biolabs). In the second step the reaction with Xrn-1 (200 μL) was carried out for 2 h at 37°C and contained digested RNA, 1 U/μL RiboLock RNase Inhibitor (Thermo Fisher Scientific), Xrn-1 buffer (New England Biolabs) and Xrn-1 (New England Biolabs), using 1 μL of enzyme per 1 μg of uncapped RNA*. The samples were then purified again with Monarch RNA Clean-Up (New England Biolabs). Finally, all RNA samples were purified using HPLC and Phenomenex Clarity® 3 μM Oligo-RP column 150 × 4.6 mm (using method: A: 100 mM TEAAc, B: 200 mM TEAAc/ACN; 10–35 % of eluent B in 32 min), precipitated at sodium salts (3M NaOAc pH 5.2, isopropanol) and dissolved in water. The micromolar concentrations of the RNA samples were determined using NanoDrop (Thermo Fisher Scientific).

### Expression and Purification of IFIT1

Human interferon-induced protein with tetratricopeptide repeats 1 (IFIT1) gene (Gene ID: 3434) was obtained from Addgene in the plasmid vector pET28a_IFIT1, dedicated to protein expression. The hIFIT1 protein (∼55 kDa) with 6x histidine at the 5’-end (His-Tag) was overexpressed in the BL21 (DE3) RIL *E. coli* (Invitrogene) procaryotic expression system in LB medium supplemented with kanamycin (30 mg/mL). Cells were grown to the optical density OD_600_∼0.7 at 37°C, then the temperature was adjusted to 18°C, expression was induced with 0.5 mM isopropyl b-D-1-thiogalactopyranoside (IPTG) and the cells were further cultured for 16 h. Bacterial cultures were harvested by centrifugation and lysed in a buffer containing 20 mM HEPES (pH 7.5), 250 mM NaCl, 20 mM imidazole, 5 mM β-mercaptoethanol (2-ME), 5% glycerol, 1mg / mL of lysozyme and mixture protease inhibitors (Aprotinin, Pepstatin, PMSF, Leupeptin). The lysate was sonicated (15 min, amplitude 50%, 15 s on/off) and centrifuged. The supernatant was then purified by immobilized metal affinity chromatography (IMAC). Protein solution was loaded on 2 × 5 mL HisTrap FF™ column (Cytiva) previously equilibrated with a buffer containing 20 mM HEPES (pH 7.5), 250 mM NaCl, 20 mM imidazole, 5 mM β-mercaptoethanol (2-ME) and 10% glycerol. The His6-IFIT1 protein was washed with a buffer containing 1 M NaCl and eluted with buffer: 20 mM HEPES (pH 7.5), 250 mM NaCl, 300 mM imidazole, 5 mM β-Mercaptoethanol and 10% glycerol. Affinity chromatography on a heparin column was then used to remove the remaining endogenous nucleic acids and some protein contaminants from the preparation (after IMAC). The protein fractions collected were diluted with 20 mM HEPES (pH 7.5), 1 mM DTT to a final salt concentration of ∼100 mM NaCl and loaded onto a 5 ml HiTrap Heparin HP ™ column (Cytiva) previously equilibrated with a buffer containing 20 mM HEPES (pH 7.5), 100 mM NaCl, 1 mM DTT (Phase A). The method was carried out with a constant increasing proportion of phase B at 1% / min from 0 to 100% (Phase B: 20 mM HEPES, 1000 mM NaCl, 1 mM DTT). The final step in protein purification was gel filtration. The eluted fractions were loaded onto a 75 pg HiLoad 26/600 ™ gel filtration column (Cytiva). Samples containing IFIT1 were centrifuged (10 min, 10 000 rcf, 4°C) to remove any aggregates, concentrated to a final concentration of 12 μM, flash frozen, and stored at -80°C in a buffer containing 20 mM HEPES (pH 7.5), 150 mM NaCl, 1 mM DTT, 10% glycerol. The nucleotide sequence of IFIT1 and MS analysis included in supplementary (Fig. S1).

### Protein Labelling

The stock solution of His6-IFIT1 (12 μM) was centrifuged (10 min, 10 000 rcf, 4°C) and diluted to 200 nM in MST buffer containing 20 mM HEPES (pH 7.5), NaCl, 0.075% Pluronic F-127. For cap analog ligands the buffer (buffer L1) contained 100 mM NaCl, whereas for capped RNA ligands – 200 mM NaCl (buffer L2). The RED-tris-NTA dye (NT-647, Nano Temper Technologies) was diluted in the same buffer to 100 nM. Protein and dye were mixed in a 1:1 volume ratio and incubated for 30 min at room temperature. Before the MST measurements, every sample of labelled protein was centrifuged to remove potential aggregates (10 min, 10 000 rcf, 4°C).

### Optimization of Conditions

Several optimization measurements were performed for labeled IFIT1 and ligands. In order to obtain stable conditions for the proper interaction between protein and ligands, three types of buffers were used: MST-buffer, 1x PBS, and 20 mM HEPES. To avoid aggregation and suppressing unspecific protein adsorption into capillary walls, we tested two different detergents: Tween-20 and Pluronic-F127 at concentrations of 0.01 - 0.1%. The use of BSA (0.1 mg/mL) instead of detergent was also explored. The effect of ionic strength was tested at three concentrations of NaCl (100 mM, 200 mM and 300 mM) in three types of buffers above. Furthermore, three different pH buffers were also tested: (pH 7, pH 7.5 and pH 8). Finally, 20 mM HEPES pH 7.5 buffer supplemented with 0.075% Pluronic F-127 and 100 mM or 200 mM NaCl (for free cap analogs and RNA ligands, respectively) was chosen. Under these conditions, the labelled protein was stable; no aggregation, no adsorption, no photobleaching, nor photoenhancement were observed, see Fig. S2.

### Microscale Thermophoresis

For all thermophoretic measurements, MST Premium coated capillaries (Nano Temper Technologies) were used. In all MST experiments, the final concentration of His-tag-labeled IFIT1 was set to 50 nM. 16-point half-log serial dilutions of ligands at different concentration ranges were prepared. The concentration ranges were adjusted for each analog to cover minimum 80% of the binding curve. The final samples were prepared by mixing an equal volume (10 μl) of 100 nM His-tag labelled IFIT1 with the ligand at a specified concentration. For RNA cap analogs, measurements were performed in buffer L1 (20 mM HEPES (pH 7.5), 100 mM NaCl, 0.075% Pluronic F-127) at ligand concentrations typically ranging from 10 mM to 305 nM. For capped RNA, measurements were performed in buffer L2 (20 mM HEPES (pH 7.5), 200 mM NaCl, 0.075% Pluronic F-127) and with ligand concentrations ranging from 20 μM to 610 fM.

Measurements of IFIT1-ligand interaction were performed on a Nano Temper Monolith™NT.115 instrument. Before MST measurement, samples were equilibrated for 15 minutes at room temperature in darkness, then loaded into capillaries, and inserted into the data collection instrument (with a temperature set at 25°C). The measurement of each ligand was performed in triplicate. The final RED-tris-NTA dye concentration of 50 nM yielded the fluorescence intensity of labelled IFIT1 around 600 counts at a LED power of 40%. The samples were measured at medium MST power with a pre-MST period of 5 s, a laser-on time period of 30 s, and a laser-off time period of 5 s.

### Data Analysis

The data (time-traces) were loaded into PALMIST v1.5.8 software and evaluated using a simple 1:1 binding model. For every measurement, the relative fluorescence value was determined by a rapid change in normalized fluorescence in phase II of thermophoresis caused by the temperature dependence of fluorescence (T-jump preset). Triplicates for given measurement points were averaged, but a weighted fitting was not used. The florescence values were normalized and a logarithmic concentration scale was applied. The plots were rendered using the GUSSI v1.4.2 programme.

## RESULTS AND DISCUSSION

### An MST-based direct binding assay enables determining the affinity of IFIT1 for long-capped RNAs

To investigate the interaction of various 5’ RNA cap analogs and capped RNAs with the IFIT1 protein in a reproducible and time-effective manner, we developed an MicroScale Thermophoresis (MST)-based direct binding assay. Microscale thermophoresis is a biophysical method relying on the measurement of two factors: directed molecule movement in a temperature gradient (thermophoresis) and change in fluorescence intensity upon changing temperature (TRIC)(Magnez, Bailly and Thuru 2022). The thermophoretic mobility of molecules is sensitive to small changes in the charge, size, and hydration shell caused by interactions with other compounds, making MST suitable for studying protein-ligand interactions across a wide range of ligand sizes. This, combined with a wide working range of measurable *K*_D_ values, straightforward optimization, and low sample consumption, led us to choosing this method. In our setup, IFIT1 fluorescently labeled with a commercially available red dye via N-terminal His-tag was used to measure ligand concentration-dependent changes in the thermophoretic mobility of the protein. IFIT1 labelling was carried out using standard methods, and the labeled protein showed good stability in aqueous buffers (Fig. S2). To enable the synthesis of variously capped mRNAs, we generated a library of 13 different cap analogs, including cap-0, cap-1, and their further methylated analogs, as well as several non-canonical RNA 5’ ends (Fig. 1). The set contained both 5’ capped RNA that were previously evaluated against IFIT as well as the unstudied ones, which allowed us to confront our newly developed method against literature data and gain new insights into RNA 5’ ends recognition by IFIT1. The workflow of the assay is shown in Fig. 2.

**Figure 2.**
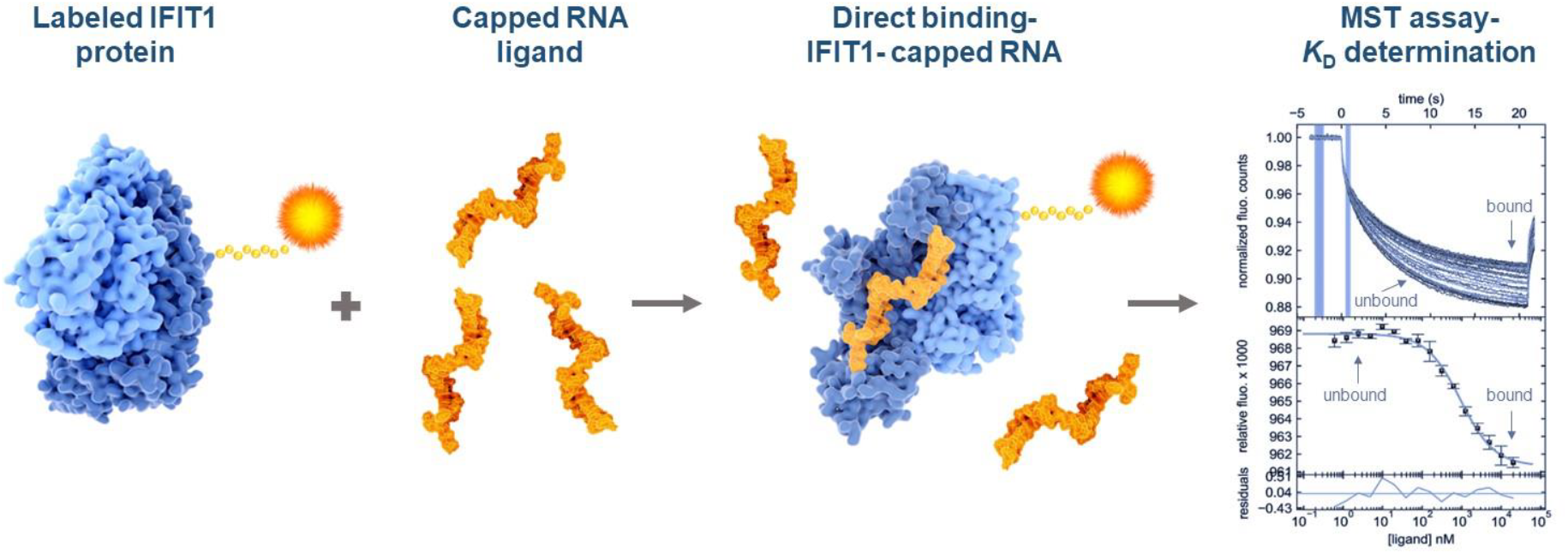
Workflow for the determination of *K*_D_ values for the IFIT1-capped RNA_70_ interaction using an MST-based binding assay. Fluorescently labeled IFIT1 was mixed with RNA ligand at 16 different concentrations, equilibrated for 15 min, followed by the analysis by MST. The observed changes in fluorescence intensity were plotted as a function of ligand concentration and a theoretical binding curve was fitted to determine the dissociation constants (*K*_D_).

To find a suitable RNA model to compare the 5′ cap variants, we first explored three different RNA sequences as IFIT1 ligands including two 35 nt RNAs differing in the secondary structure and a 70 nt fragment of human β-globin protein. For each sequence variant, RNAs capped with m^7^GpppApG (cap-0) or m^7^GpppA_m_pG (cap-1) were synthesized by an *in vitro* transcription (IVT) reaction. To ensure high purity of the analysed RNA samples, uncapped RNA molecules were removed by enzymatic digestion and the remaining material was purified by RP-HPLC. Since the preliminary analyses indicated significant differences in the affinity of RNA ligands for IFIT1 compared to the corresponding trinucleotide cap analogs, the ionic strength of the buffers for both types of ligands were differentiated to move the *K*_D_ values toward the higher end of the range (Fig. S3, Table 1A). In each case, the MST analysis (Fig. 3, Table 1) showed binding curves that were consistent with a 1:1 binding model (Fig. 3).

**Table 1 A.** Binding affinities of IFIT1 determined by MST for cap-0 and cap-1 structures depending on.

| Cap structure | $K_D$ (C.I.) / [nM] | | | |
| --- | --- | --- | --- | --- |
|  | RNA I [a],[c] | RNA II [a],[c] | RNA III [a],[c] | Trinucleotide [b] |
| m<sup>7</sup>GpppApG | 8 (1–19) | 15 (8–25) | 100 (60–170) | 100 (60–150) |
| m<sup>7</sup>GpppA<sub>m</sub>pG | 34 (19–59) | 60 (42–84) | 520 (350–770) | 380 (190–610) ·10<sup>3</sup> |

**Table 1 B.** The quotient of the *K*_*D*_ values of the cap-1 and cap-0 for all tested sequence variants.

| $K_D$ cap-1/ $K_D$ cap-0 | RNA I [a] | RNA II [a] | RNA III [a] | Trinucleotide [b] |
| --- | --- | --- | --- | --- |
| cap-1 / cap-0 | 4.2 | 4.0 | 5.2 | 3800 |

[a] conditions: 20 mM HEPES (pH 7.5), 200 mM NaCl, 0.075% Pluronic F-127, mean values from triplicate experiments with confidence intervals (C.I.).
[b] conditions: 20 mM HEPES (pH 7.5), 100 mM NaCl, 0.075% Pluronic F-127, mean values from triplicate experiments with confidence intervals (C.I.).
[c] RNA sequences are shown in Figure 3 and in the Experimental Section.

**Figure 3.**
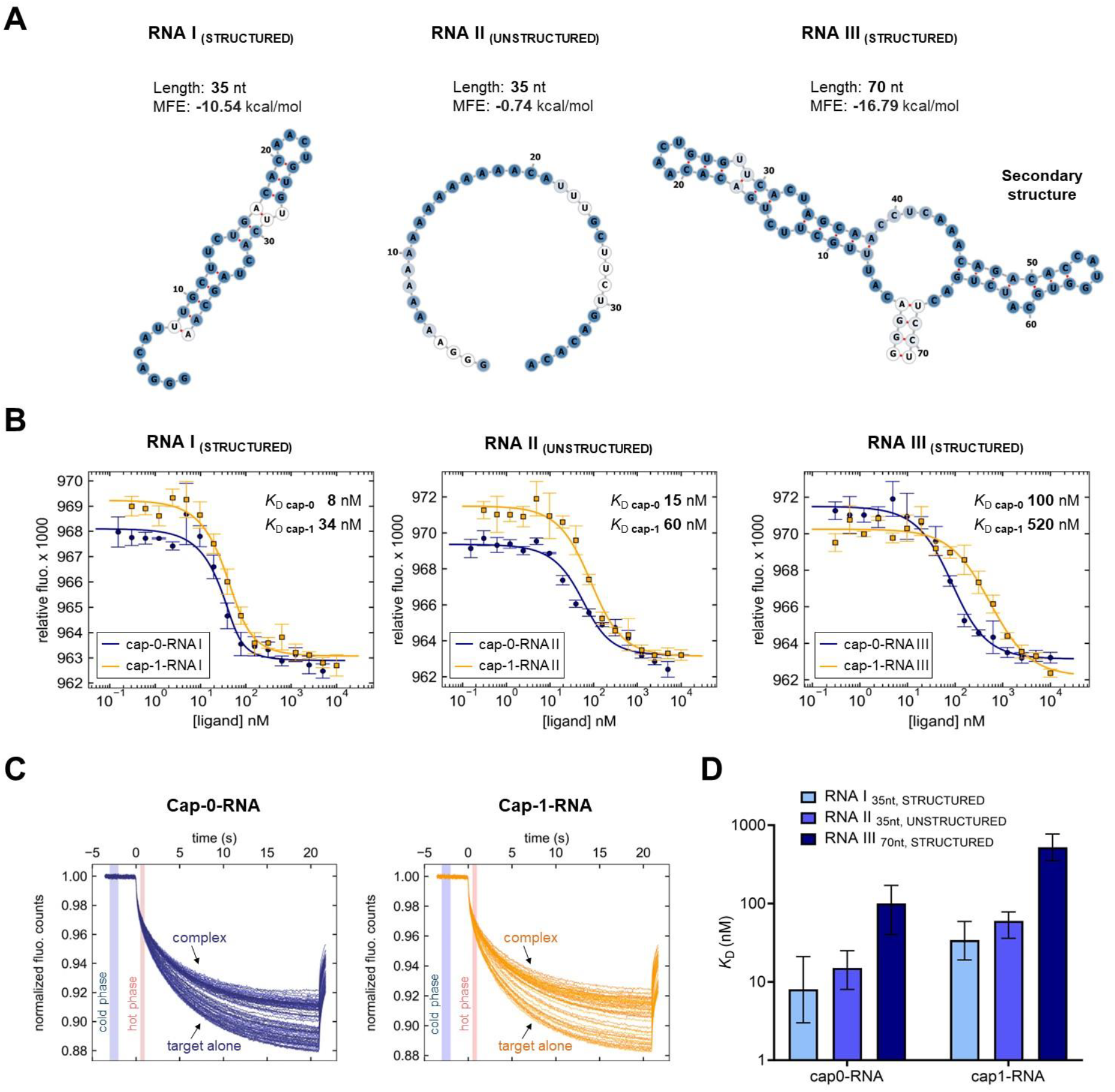
Determining *K*_D_ values for three different sequence variants of cap-0 and cap-1 RNA using the MST method. A) Predicted secondary structures of the analyzed RNA sequences (model Vienna RNAfold) and their MFE (minimum free energies). B) The determined MST-binding curves from triplicate experiments (data points represent mean values ± SD. Binding curves are best fits for 1:1 binding model. C) Representative MST traces for two of the curves shown in B (RNAIII). Vertical marks (blue and red highlights) indicate cold and hot regions that were used for further analysis (T_jump_). D) Comparison of the determined *K*_D_ values (mean values from triplicate experiments with confidence intervals).

**Figure 4.**
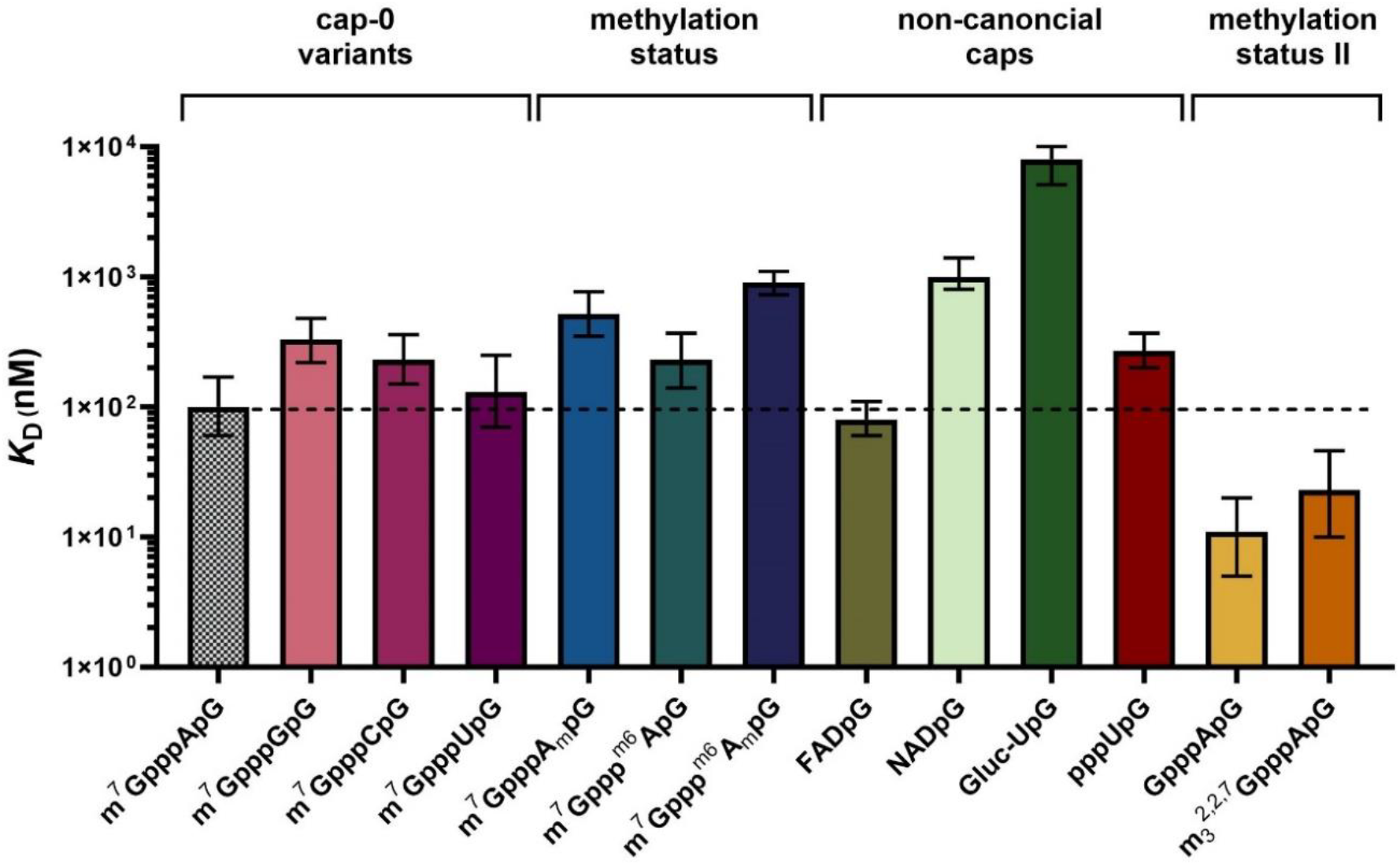
Comparison of *K*_D_ values for all measured capped 70 nt RNAs. Data shown are mean values from minimum three replicates with confidence intervals.

The RNA I sequence forming a hairpin structure (Fig. 3A), bound to IFIT1 with *K*_D_ values of 8 nM for cap-0 and 34 nM for cap-1. The RNA II (unstructured, 35 nt) that was designed to avoid formation of any secondary structures, bound to IFIT1 with the slightly higher *K*_D_ values: 15 nM for cap-0- and 60 nM for cap-1. For the third sequence variant, RNA III (structured, 70 nt), we observed notably weaker binding than for shorter RNAs, with *K*_D_ values of 100 nM for cap-0 and 520 nM for cap-1. In each case, the affinity of IFIT1 for RNAs with cap-0 structure was 4–5 fold higher than for RNAs with cap-1, which is consistent with previous results, and indirectly confirms that the differentiation of 5’-terminal structures by IFIT1 is RNA sequence independent.

The differences in the affinity exhibited by diverse RNA sequences may be attributed to electrostatic interactions between the RNA body and protein surface. RNA III sequence provided the relatively highest difference between cap-0- and cap-1-RNA (Table 1B), whereas RNA I and II afforded *K*_D_ values closer to the lower limit of the working range of our MST method (∼1**–**10 nM), which could impede the evaluation of high-affinity ligands. Hence, we reasoned that RNA III is the most suitable model for evaluating the influence of differently modified cap structures on RNA affinity for IFIT1.

### Development of a direct binding assay for the evaluation of the IFIT1-capped RNA_70_

Next, we prepared a set of eleven additional variants of RNA III differing in the 5’ terminal modifications and determined their binding affinities for IFIT1 (Table 2). These variants included four additional cap-0 variants differing in the first transcribed nucleobase (G, C, U, m^6^A), cap-1 variant containing m^6^A_m_ mark, and six non-canonical 5’ ends including G-cap, NAD, FAD, TMG, UDP-glucose, and UTP.

**Table 2.** The *K*_D_ values of binding between IFIT1 and 5’ RNA cap analogs or 5’ capped RNAs.

| 5' RNA cap analog | Capped RNA<sub>70</sub><sup>[a],[c]</sup><br>$K_D$ [nM] | Capped RNA<sub>70</sub><sup>[d]</sup><br>C.I. [nM] | Trinucleotide cap<sup>[b]</sup><br>$K_D$ [nM] | Trinucleotide cap<sup>[d]</sup><br>C.I. [nM] |
| --- | --- | --- | --- | --- |
| m<sup>7</sup>GpppApG | 100 | (60, 170) | 100 | (60, 150) |
| m<sup>7</sup>GpppAmpG | 520 | (350, 770) | $380 \cdot 10^3$ | $(190, 610) \cdot 10^3$ |
| m<sup>7</sup>GpppCpG | 230 | (150, 360) | $13 \cdot 10^3$ | $(10, 15) \cdot 10^3$ |
| m<sup>7</sup>GpppGpG | 330 | (220, 480) | $23 \cdot 10^3$ | $(14, 39) \cdot 10^3$ |
| m<sup>7</sup>GpppUpG | 130 | (70, 250) | $390 \cdot 10^3$ | $(330, 440) \cdot 10^3$ |
| m<sup>7</sup>Gpppm<sup>6</sup>ApG | 230 | (140, 370) | $50 \cdot 10^3$ | $(20, 100) \cdot 10^3$ |
| m<sup>7</sup>Gpppm<sup>6</sup>AmpG | 900 | (730, 1100) | $380 \cdot 10^4$ | $(210, 780) \cdot 10^4$ |
| GpppApG | 11 | (5, 20) | $370 \cdot 10^4$ | $(210, 660) \cdot 10^4$ |
| NADpG | 1000 | (800, 1400) | $430 \cdot 10^6$ | $(200, 540) \cdot 10^6$ |
| m<sub>3</sub><sup>2,2,7</sup>GpppApG | 23 | (10, 46) | 30 | (19, 47) |
| FADpG | 80 | (60, 110) | $21 \cdot 10^3$ | $(13, 32) \cdot 10^3$ |
| pppUpG | 270 | (200, 370) | $26 \cdot 10^6$ | $(14, 112) \cdot 10^6$ |
| Glc-ppUpG | 8 200 | (6 200, 11 100) | n/b<sup>[e]</sup> | n/b<sup>[e]</sup> |

[a] conditions: 20 mM HEPES (pH 7.5), 200 mM NaCl, 0.075% Pluronic F-127, mean values from triplicate experiments.
[b] conditions: 20 mM HEPES (pH 7.5), 100 mM NaCl, 0.075% Pluronic F-127, mean values from triplicate experiments.
[c] RNA sequences is RNA III shown in Figure 3 and given in the Experimental Section.
[d] C.I. (Confidence Interval) values estimated automatically by PALMIST- EST procedure (Error Surface Projection). Confidence level set at 68.3% (equivalent to $\pm 1\sigma$ in Gaussian bell curve).
[e] n/b- no binding was observed up to concentration of 40 mM ligand.

The comparison of affinities for RNAs carrying different cap-0 variants (A, G, C, U, m^6^A) revealed a mild preference for A and U at the position of the first transcribed nucleotide (Table 2). The affinities were highest for m^7^GpppApG, followed by 1.3-fold weaker interaction for m^7^GpppUpG-RNA, 2.3-fold weaker for m^7^GpppCpG and ending with 3.3-fold weaker for m^7^GpppGpG. Notably, the lowest affinity obtained for the cap-0 RNA variant (m^7^GpppGpG-RNA) was only 1.6-fold higher than that of cap-1 carrying adenine (m^7^GpppA_m_pG-RNA; Table 2). The methylation of cap-0 at the N6 position of adenosine (m^7^Gppp^m6^ApG-RNA) lowered the affinity of RNA for IFIT1, but not as much as methylation at the 2’-*O* position (the *K*_D_ value of m^7^Gppp^m6^ApG-RNA was 2.3-fold higher than that of m^7^GpppApG-RNA but 2.2-fold lower than that for m^7^GpppA_m_pG-RNA). Notably, the effects of these two methylations were additive since the RNA probe carrying m^7^Gpppm^6^A_m_pG cap had 9-fold lower affinity compared to cap-0 RNA ligand. This suggests that the N6-methylation of the 5’-terminal adenosine constitutes an additional layer of protection from RNA sequestering by IFIT1.

The investigated non-canonical caps had a very variable effect on IFIT1-RNA interaction. The *K*_D_ values determined for non-canonically capped RNAs showed broad affinity spectrum that extended over three orders of magnitude (Table 2). FADpG-RNA bound to IFIT1 with affinity comparable to cap-0 RNA (*K*_D_ 80 nM; C.I. 60–110 nM). GpppApG-RNA, an intermediate during cap biosynthesis (Shannon et al. 2022), was bound by IFIT1 even more avidly than cap-0-RNA (*K*_D_ 11 nM). Surprisingly, the hypermethylation of the m^7^G moiety leading to the so-called trimethylguanosine cap (m_3_ ^2,2,7^GpppApG-RNA) also resulted in a significant increase in the affinity for IFIT1 (*K*_D_ 23 nM). NADpG-RNA had a significantly lower affinity (*K*_D_ 1.0 μM), and Glucose-ppUpG-RNA was a very weak binder (*K*_D_ 8.2 μM), much weaker than the corresponding triphosphate RNA (*K*_D_ 270 nM). Overall, these results may indicate that IFIT1 is capable of targeting more RNA 5’ terminal structural patterns than previously anticipated, and that some non-canonical 5’ end modifications may protect RNA from binding by IFIT1.

Finally, we also wanted to test if the use of short, trinucleotide cap analogs as IFIT1 ligands would provide meaningful data on the relative affinities of different cap variants. This could significantly reduce the time and the total cost of the analysis. The *K*_D_ values determined for m^7^GpppApG (cap-0) and m^7^GpppA_m_pG (cap-1) suggested that IFIT is able to differentiate caps even more strongly if the RNA body is not present (Table 1, Table 2). However, a back-to-back comparison of all *K*_D_ values determined for the capped RNA ligands and free caps (Table 2) suggests that the measurements for free cap analogs do not provide reliable quantitative data, albeit to some extent are in qualitative agreement with the data obtained for RNA ligands. High binding affinity was observed for some particularly strongly interacting RNA 5’ ends, even for “free” caps (m_3_^7^GpppApG, m_3_ ^2,2,7^GpppApG). This may suggest that these structures specifically interact with IFIT1 and are particularly significant as its cellular targets.

## CONCLUSIONS

The continuous increase in the number of potential applications of RNA molecules highlights the need for the development of better methods to understand the structure-function relationships and biological effects for various RNA modifications. We have developed a robust and reproducible assay to evaluate the affinity of differently capped RNAs for IFIT1. The assay, which relies on measuring microscale thermophoresis of fluorescently labelled protein in the presence of increasing ligand concentrations, enables more efficient analysis of IFIT1 RNA interactions compared to previously developed methods due to its low sample consumption, simple workflow and easily quantifiable results. We have demonstrated that the assay is suitable for analysis of 35 and 70 nt long RNA ligands, but there are no technical obstacles to the analysis of either shorter or longer RNAs. We also showed that the analysis of small molecule cap analogs is technically possible using the MST assay, yet it does not deliver results that can be easily translated into properties of longer RNAs. Still, some qualitative trends in the structure-affinity-relationship observed for RNA ligands are noticeable also for trinucleotide cap analogs, which suggests that the assay may be applicable for discovering high-affinity small molecule ligands of IFIT1 if desired.

We used the developed assay to back-to-back compare a set of thirteen RNA ligands carrying different natural canonical and non-canonical cap structures. To our knowledge, this is the first such a broad comparison of differently capped RNAs in the context of interactions with IFIT1, and many cap modification studies here have not been previously investigated. Starting with the reference molecules, the affinity for cap-0 was about 5 times higher than for the cap-1 molecule. All four tested cap-0 RNAs showed comparably strong binding to IFIT1 in the *K*_D_ range of 100–300 nM. Comparing unmethylated ApG (cap-0) and ^m6^ApG RNA ligands we saw a decrease in affinity for the methylated analogue, and the effect was even greater when combined with 2′-*O*-methylation to produce ^m6^A_m_. This indicates a trend of weakening the interaction of the IFIT1 protein with 5’ RNA structures that have additional methylations on the first transcribed nucleotide. For the m^7^G modifications, both the deletion of methylation from m^7^G (GpppApG) and hypermethylation (TMGpG) resulted in increased affinity towards IFIT1. Although both these structures are formed during biosynthesis or maturation of some of endogenous RNAs (Terns and Dahlberg 1994, Wurth et al. 2014), their exceptionally high affinity of IFIT1 for this hypermethylated structure implies that they may be another class of cellular IFIT1 targets.

The analysis of non-canonical cap structures revealed very high divergence in affinity. The FAD-capped RNA ligand strongly binds to IFIT1, while NAD-capped RNA binds more than 10 times weaker, and the UDP-glucose-capped RNA is hardly recognized by IFIT1. The UDP-glucose moiety has a clearly “protective” effect against IFIT1, as it has an affinity lower than the corresponding uncapped RNA.

Interestingly, all structures that have been identified in this study as tightly binding to IFIT1, have been previously somehow linked to viral RNA. Since mRNA is capped in the nucleus, GpppApG-RNA is normally absent in the cytoplasm but is formed there during some viral RNA capping pathways. Besides its presence on endogenous RNAs, trimethylguanosine cap has also been identified on some HIV-associated RNAs (Boris-Lawrie et al. 2022, Qiu, Shuman and Schwer 2011). The 5’-terminal FAD cap has been found to modify about 75% of HCV RNAs (Sherwood et al. 2023). The fact that our in vitro assay revealed comparably high or even higher affinity of these three molecular patterns to cap-0 RNA may suggest that they are also targeted by IFIT1. The biological relevance (if exists) of non-canonical modifications that had significant protective effects against IFIT1, with UDP-glucose being the most striking example, needs to be verified, but it may also guide future design of non-coding RNAs with therapeutic function.

The use of mRNA in therapies offers numerous advantages over conventional methods, as evidenced by the rapid development of mRNA technology in recent years (Yao, Xie and Xia 2024, Huang et al. 2022, Xu et al. 2020). However, the administration of exogenous mRNA to patients carries the risk of triggering an innate immune response against foreign RNA (Schlee and Hartmann 2016). The innate immune response executed by IFIT1 can efficiently and selectively influence the translation by shutting down selected viral and cellular mRNAs and inhibiting the amplification of foreign genetic material. However, these same mechanisms may reduce the efficacy of mRNA-based therapeutics (Andrejeva et al. 2013).The influencing factor in the occurrence of such adverse effects is the recognition of therapeutic mRNA molecules by IFIT1 (Abbas et al. 2017, Pingale, Kanade and Karpe 2019, Russ et al. 2022). The incorporation of cap-1 in therapeutic mRNAs is anticipated to mitigate the risk of an adverse immune response while concurrently reducing susceptibility to translational inhibition mediated by IFIT1 (Johnson et al. 2018, Fleith et al. 2018, Pinto et al. 2015). Recent discoveries increasingly underscore the importance of chemical modifications of RNA in modulating the innate immune response (Tong et al. 2022). The use of improved mRNA molecules in therapy has the potential to enhance the translation of the target protein, expediting therapeutic effects (Baptista et al. 2021). We hypothesize that the design of chemically modified mRNA cap structures that can evade sequestering IFIT1 may additionally benefit the therapeutic mRNA field. We hope that the assay reported will become an essential tool for verifying this hypothesis as well.

## SUPPLEMENTARY DATA

Supplementary Data are available at.

## AUTHOR CONTRIBUTIONS

TS and JK designed the study. JJ and JK supervised the study. TS performed experiments. AD, KG, KZ and MW provided resources. TS and JK wrote the first draft of the manuscript. The manuscript was written through contributions of all authors. All authors have given approval to the final version of the manuscript.

